# Targeted metagenomics reveals pangenomic diversity of the nitroplast (UCYN-A) and its algal host plastid

**DOI:** 10.1101/2024.06.19.599377

**Authors:** Ella Joy H. Kantor, Brent M. Robicheau, Jennifer Tolman, John M. Archibald, Julie LaRoche

**Author notes:** The authors declare that they have no competing interests.

## Abstract

UCYN-A (*Cand.* Atelocyanobacterium thalassa) has recently been recognized as a globally-distributed, early stage, nitrogen-fixing organelle (the ‘nitroplast’) of cyanobacterial origin present in select species of haptophyte algae (e.g., *Braarudosphaera bigelowii*). Although the nitroplast was recognized as the UCYN-A2 sublineage, it is yet to be confirmed in other sublineages of the algal/UCYN-A complex. We used water samples collected from Halifax Harbour (Bedford Basin, Nova Scotia, Canada) and the offshore Scotian Shelf to further our understanding of *B. bigelowii* and UCYN-A in the coastal Northwest Atlantic. Sequencing data revealed UCYN-A-associated haptophyte signatures and yielded near-complete metagenome-assembled genomes (MAGs) for UCYN-A1, UCYN-A4, and the plastid of the A4-associated haptophyte. Comparative genomics provided new insights into the pangenome of UCYN-A. The UCYN-A4 MAG is the first genome sequenced from this sublineage and shares ∼85% identity with the UCYN-A2 nitroplast. Genes missing in the reduced genome of the nitroplast were also missing in the A4 MAG supporting its likely classification as a nitroplast as well. The UCYN-A1 MAG was found to be nearly 100% identical to the reference genome despite coming from different ocean basins. Time-series data paired with the recurrence of specific microbes in enrichment cultures gave insight into the microbes that frequently co-occur with the algal/UCYN-A complex (e.g., *Pelagibacter ubique*). Overall, our study expands knowledge of UCYN-A and its host across major ocean basins and investigates their co-occurring microbes in the coastal Northwest Atlantic (NWA), thereby facilitating future studies on the underpinnings of haptophyte-associated diazotrophy in the sea.

## Introduction

Diazotrophic microbes play a key role in the marine nitrogen cycle by converting biologically unavailable atmospheric dinitrogen (N_2_) into ammonia, which can then be assimilated into organic nitrogen (1). N_2_ fixation is accomplished by the multi-subunit nitrogenase enzyme, which is inactivated by oxygen and has high iron requirements (2,3). Marine diazotrophs are typically free-living or symbiotic cyanobacteria, although non-cyanobacterial diazotrophs are also known, with various groups having evolved different strategies for overcoming oxygen constraints (4). The main gene used to track diazotrophs is *nifH,* which codes for a subunit of the nitrogenase enzyme (5,6). *NifH* signatures throughout the world’s oceans have led to the discovery of widely distributed unicellular diazotrophs, which are now known to contribute significantly to global N_2_ fixation (7,8). The *nifH* gene of *Candidatus* Atelocyanobacterium thalassa (or UCYN-A (9)), a cyanobacterial sequence, exhibited a cosmopolitan distribution and was associated with high N_2_ fixation rates (6,10,11). The endosymbiotic lifestyle of UCYN-A was hypothesized early on due to its highly reduced genome and tight association with a haptophyte alga, with which it exchanges fixed nitrogen in return for plastid-derived fixed carbon (9,12). However, recent evidence suggests that UCYN-A is in fact a fully integrated nitrogen-fixing organelle, dubbed a “nitroplast” (13).

Up to eight UCYN-A sublineages or phylotypes—called A1 to A8—have been proposed through *nifH* metabarcoding and have also been considered ecotypes (14–16). In a recent study of UCYN-A in the coastal Northwest Atlantic, Robicheau et al. (17) described the co-occurrence of sublineages A1-A4. Network analysis of the phytoplankton temporally associated with UCYN-A in this region found a strong copresence of UCYN-A2 *nifH* with the *16S* rRNA plastid signature of the known haptophyte host *Braarudosphaera bigelowii*, as well as a copresence between UCYN-A1 and another haptophyte, *Chrysochromulina* sp., a potential host for this latter ecotype (17,18). Seven UCYN-A genomes are currently publicly available, three from the A1 sublineage and four from A2 (Table S1). Three putative A2-associated *B. bigelowii* partial plastid sequences have also been published (13).

Evidence that UCYN-A is an evolving organelle was obtained using the *B. bigelowii*-associated A2 ecotype. Coale et al. (13) demonstrated genome reduction, synchronized division with the host, and the presence of host nuclear encoded proteins in the UCYN-A proteome. We use the term UCYN-A to describe the *Candidatus* Atelocyanobacterium thalassa derived ecotypes and genomes (i.e. A1, A2, etc.) to avoid confusion with older publications that historically use this terminology/notation, however, UCYN-A2 is the newly identified nitroplast of *B. bigelowii*.

Here, we used enrichment culturing to select for diazotrophs within a mixed microbial community then identified samples that contained higher proportions of UCYN-A. Amplicon and metagenomic sequencing were used to characterize microbes in the UCYN-A-containing cultures and to generate metagenome-assembled genomes (MAGs). We present a UCYN-A4 ecotype MAG and its algal-associated plastid sequence, as well as a pangenome analysis of available UCYN-A genomes. We further analyze microbes co-occurring with UCYN-A both in enrichment cultures and in the natural communities of the coastal Northwest Atlantic using a weekly multi-year oceanographic time series.

## Materials and Methods

### Water collection, enrichment culturing, and cell sorting

Seawater was collected via Niskin bottles from the offshore Scotian Shelf (at 20m) in 2021 [Station HL7 during 2021 MORI Atlantic Condor Expedition; 42° 50’ 00” N, 61° 43’ 00” W] and weekly from the inshore Bedford Basin (at 5m) during late summer/early fall of 2018 and 2020 [at 44° 41’ 37“ N, 63° 38’ 25” W; Halifax, N.S., Canada]. The sampling period targeted UCYN-A in this region (17). Samples were enriched with iron (FeCl_3_), phosphate (NaH_2_PO_4_), and vitamins (Biotin, Cobalamin, Thiamine-HCl; Table S2) to select for diazotrophs under laboratory conditions (see (19,20)). Incubations occurred in polystyrene tissue culturing flasks (Greiner Bio-One, Austria) at 15°C and 12h light/dark cycles for at least two weeks to allow for the stabilization of the community prior to further experimentation. Following this initial incubation, cultures were screened using quantitative-PCR (qPCR) *nifH* assays for UCYN-A1 or A2/3/4 using DNA extracted from filtered cells (17,21)–noting that the A2 assay used is cross-reactive to A3/A4 (17,22). Samples with high UCYN-A *nifH* counts were retained for further analysis.

One sample from the Scotian Shelf (Shelf 1) and one sample from the Bedford Basin (Basin 1) had a high abundance of UCYN-A1 directly following initial nutrient amendment and multi-week incubation; cells from these samples were directly used for DNA extractions and sequencing without cell sorting. Other samples containing UCYN-A2/3/4 were subjected to fluorescence-activated cell sorting (FACS) using a BD Influx Cell Sorter to further concentrate cells containing UCYN-A for additional culturing/collection. Sorted populations were screened using the qPCR assays used above (17,21). For the other Bedford Basin sample (Basin 2) we sorted 2,000 cells from a cytogram population attributed to UCYN-A into the 0.2μm filter-sterilized culture medium. Sorted cells were immediately transferred and grown for another 8.5 weeks in ∼10mL of 0.2μm-filtered Bedford Basin seawater that had been archived from the same day as the original sample. This seawater was further enriched with Fe, PO_4_, and vitamins after 0 weeks, 2 weeks, and 4 weeks of secondary incubation (same concentrations as in Table S2). For the Scotian Shelf sorted sample (Shelf 2) that contained mainly UCYN-A2/3/4, we did not have to re-culture sorted cells to achieve a high biomass for downstream molecular work. Instead, we used ∼28,000 sorted cells attributed to a UCYN-A cytogram population (qPCR-screened) for direct DNA extraction and sequencing. Liquid enrichment cultures were harvested for molecular work, by filtration onto 0.2μm polycarbonate Isopore filters to collect biomass from 30-50mL of culture; alternatively, we used sorted cells directly.

In addition to seawater collected especially for culturing work, weekly Bedford Basin time-series seawater was used for microbial community composition analyses via *16S* rRNA metabarcoding. We therefore use the term “UCYN-A enrichment cultures” throughout to specify the difference between datasets that were derived from longer-term incubations and/or cell sorting, and those of the natural *in-situ* community. Methods used in collecting Bedford Basin time-series water samples for *in-situ* work have been described elsewhere (23).

### DNA extractions and sequencing

DNA was extracted using a DNeasy Plant Mini kit and according to the manufacturer’s instructions (Qiagen, Germany) along with a modified lysis procedure (24). Where specified, DNA was sent for metagenomic sequencing on an Illumina NextSeq instrument (Table S3) and for metabarcoding of the 18S V4 and 16S V6-V8 rRNA regions on an Illumina MiSeq at the Integrated Microbiome Resource at Dalhousie University (Halifax, N.S., Canada). Downstream analysis on amplicon sequencing data was done using the same methods and pipelines as in Robicheau et al. (23).

### Data Analyses

Metagenomic sequences were processed into bins and metagenome assembled genomes (MAGs) using the workflow in Anvi’o v7.1 (see: merenlab.org/2016/06/22/anvio-tutorial-v2/; last accessed: 31-May-2024) as eight samples in one set (25). Commands used to process raw reads into a contigs database with open reading frames (ORFs) identified are given in Supplementary Methods S1. The final contigs database that includes ORFs was populated with annotations, first with the identification of single-copy core genes (SCGs) using Hidden Markov Models by HMMR (26). SCGs were taxonomically classified using The Genome Taxonomy Database (GTDB) (27). The gene calls were classified taxonomically using Kaiju (28) and functionally annotated using NCBI’s COG, Pfam protein family database, and the KOfam database of KEGG orthologs (29–31). Sorting and indexing of the BAM files from the mapping was done using SAMtools (32). Then the mapping results were profiled using ‘anvi-profile’, which characterizes properties of every contig in a sample such as coverage and single nucleotide variants into a profile database; the profile databases for each sample were merged into one. Automatic binning was done initially using METABAT2 (33) then manually adjusted using the Anvi’o interactive interface to separate bins with high redundancy following the online tutorial (merenlab.org/2015/05/11/anvi-refine/; last accessed: 31-May-2024) (34). Bins were identified as MAGs if they met the threshold of over 70% completion or over 2 megabase pairs (Mbp) in size, and below 10% redundancy.

The pangenome was analyzed using the Anvi’o pangenomics workflow (merenlab.org/2016/11/08/pangenomics-v2/; last accessed: 31-May-2024; (35). UCYN-A genomes used include the two UCYN-A MAGs generated herein via Anvi’o and seven other published genomes which we refer to by the names given in table S1. Prior genomes include the complete reference genome for A1 of Tripp et al. (36), and for the A2 (the nitroplast), Suzuki et al. (37). The UCYN-A3 genome is partial, so only 16S rRNA and *nifH* genes were used for this analysis (38). Genomes were first uploaded into Anvi’o v7.1 and a contigs database was created for each genome and populated using the annotation steps given above and in Supplementary Methods S1, excluding Kaiju taxonomy. The commands used to generate the pangenome database, the calculation of average nucleotide identity (ANI) and pangenome visualizations are given in Supplementary Methods S2.

A MAG for the plastid genome of *B. bigelowii* was assembled using only the sequences from the Shelf 2 sample. This sample was run through the same Anvi’o metagenomics workflow described earlier except all binning was done manually. The published plastid genome of *Chrysochromulina parva* was used as a reference and the bins containing GC content similar to it (∼30%) were pulled for further examination. Contigs with high BLAST similarity to the *Chrysochromulina parva* reference were used as scaffolds on which to carry out further assembly and mapping of the raw reads using Geneious Prime 2023.2.1 (https://www.geneious.com). Plastid genome annotations shown herein and synteny were determined using Supplementary Methods S3.

16S rRNA amplicon data were processed according to Supplementary Methods S4. Geneious Prime 2023.2.1 (https://www.geneious.com) was used to locally align (via BLAST (39)) the *nifH* genes of the MAGs against UCYN-A *nifH* ASVs in Robicheau et al. (17) and oligotypes in Turk-Kubo et al. (14), with the goal of assigning UCYN-A genomes to specific sublineages. The N_2_ fixation gene region was identified using gene annotations and the Nif operon described by Zehr et al. (40). Clustal Omega v1.2.3 (41) was used to align the nitrogenase coding regions across genomes and to align 16S and 18S rRNA sequences from the metagenomic and amplicon sequencing data.

Additional annotation was done using the RAST annotation service (42–44). Proksee was used for visualization of genomes and BLAST results (45). MAUVE v1.1.3 was used to align MAGs to reference genomes to compare genes (46). CheckM2 v1.0.1 (47) was used to calculate genome completeness and contamination in addition to the tool built into Anvi’o. Completeness predictions for all nine genomes were done using the ‘checkm2 predict’ command and used the specific neural network model.

## Results

### Community Composition of Enrichment Cultures

Though not axenic, UCYN-A enrichment cultures were reduced in complexity and had a relatively higher proportion of UCYN-A either from shifting the community towards diazotrophs and/or due to the sorting procedures that directly captured UCYN-A containing cells. Metabarcoding of enrichment cultures yielded two UCYN-A ASVs [ASV022 + ASV023] across the four cultures, as well as two ASVs for *Braarudosphaera bigelowii* plastids [ASV020 + ASV021] (Figure 1A-D). The cell-sorted Shelf 2 sample had the lowest total ASV richness; this sample almost exclusively consisted of ASVs from the *B. bigelowii* plastids and UCYN-A4 (Figure 1D). In other samples some taxa occurred at comparably high proportions to the *B. bigelowii*/UCYN-A ASVs, indicating they were co-enriched during *ex situ* culturing (Figure 1A-C). Such ASVs belonged to *Pelagibacter ubique, Polaribacter* sp.*, Synechococcus* sp., and in the Basin samples, an unknown Saprospiraceae (Figure 1A, B). For the enriched (but not sorted) Shelf 1 sample, *Roseobacter* sp., *Thalassolituus* sp., *Alteromonas* sp.*, Pseudoalteromonas* sp., and *Pelagibacter ubique* were co-enriched (Figure 1C). These additional ASVs were also detected in the 4-year weekly Bedford Basin 16S rRNA time-series dataset (Figure 1E). Some of the taxa occur in the Bedford Basin throughout the year irrespective of UCYN-A (e.g., ASV075 *Abyssibacter profundi*), whereas for others (e.g., *Roseobacter* sp.) their seasonal presence roughly coincides with that of *B. bigelowii*/UCYN-A (Figure 1E). In general, the most abundant microbes in the enrichment cultures also had high overall relative abundances in the Bedford Basin (Figure 1E). Although some of the broader taxonomy for the total set of ASVs that co-enriched overlaps across samples (for those ASVs with >0.1% relative abundance), there was very little overlap at the ASV level between taxa co-enriched from the Shelf samples and the Basin samples (Figure S1, S4).

**Figure 1.**
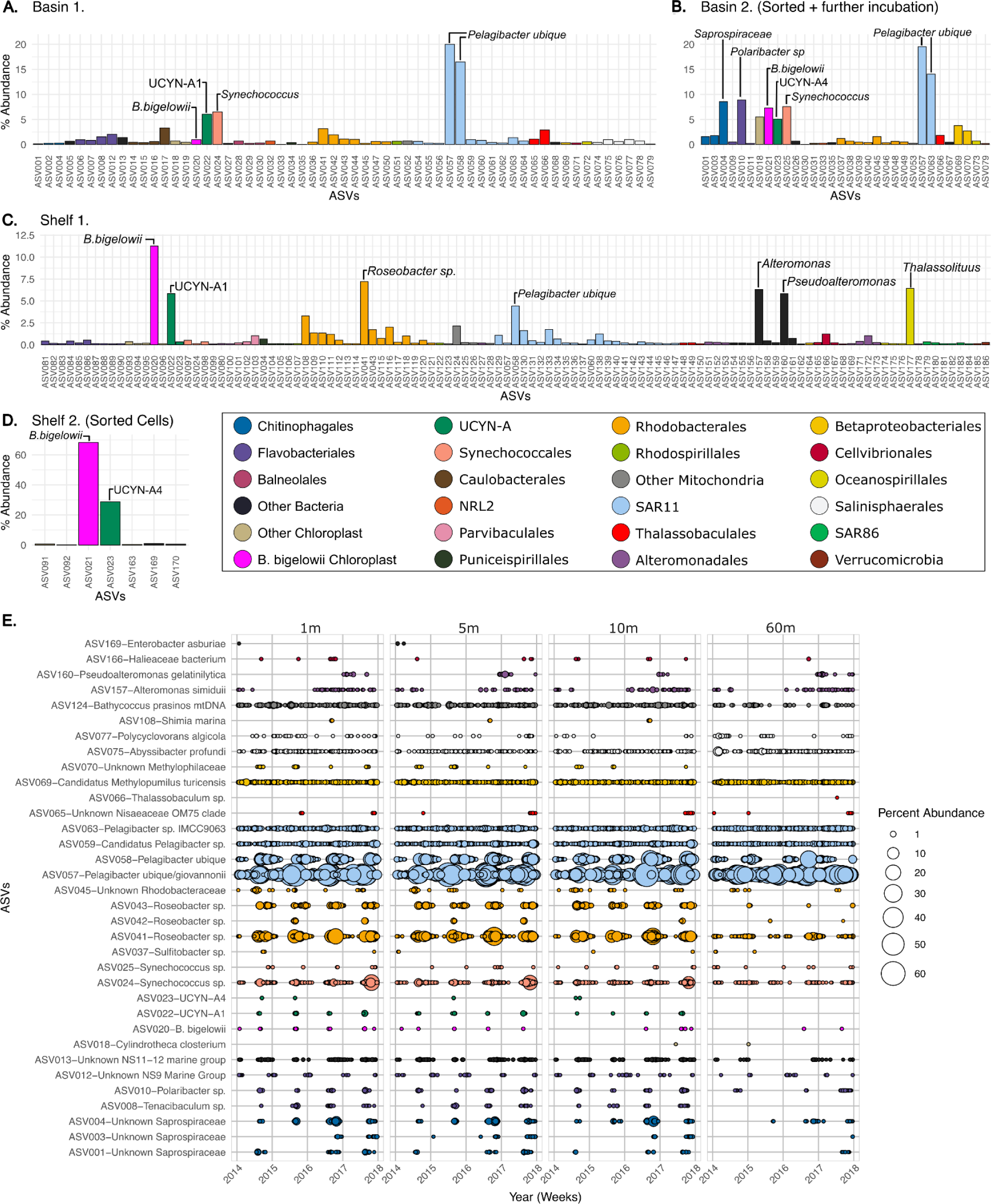
Community characterization of enrichment cultures based on relative abundance of ASVs. The results of the 16S amplicon sequencing, ASVs with ≥0.1% relative abundance are plotted for each enrichment culture used in the analyses (A-D). Panel E. uses 16S data from the Bedford Basin weekly time series to show percent abundance in the Bedford Basin of ASVs with ≥1% abundance in the enrichment cultures for each depth from the years 2014-2018.

Since they had reduced complexity, the Basin 2 and Shelf 2 samples were sequenced for 18S rRNA to further refine any putative haptophyte signatures associated with UCYN-A (see Table S5A). Fifteen 18S rRNA ASVs were only in Basin 2 and four were only in Shelf 2; the one ASV in both cultures was taxonomically classified as *Braarudosphaera bigelowii* and was the most abundant 18S rRNA ASV in the Shelf 2 sample and the second most abundant in Basin 2 after a *Cylindrotheca closterium* ASV (Table S5A). This *B. bigelowii* 18S rRNA ASV is identical to an aligned portion of 18S rRNA of Shukutsu22 isolate (AB847971.1), as well as to isolates Shukutsu27, Shukutsu19, and TP05-6-b (AB847972.1, AB847970.1, AB250785.1; Table S5B). These matching sequences are all classified as genotype I and derived from the Northwest Pacific Ocean (48,49), highlighting that for the gene region in question, the nucleotide similarity can be identical across Pacific and Atlantic ocean basins.

### MAGs of UCYN-A from the coastal Northwest Atlantic

104 bins were assembled from enrichment culture metagenomic data, of which 35 were classified as MAGs with >70% completion or >2 Mbp and redundancy <10%. Two MAGs, MAG_00024 and MAG_00029, belonged to UCYN-A genomes based on SCG taxonomy identification. MAG_00029 has seven contigs and is mostly composed of reads from Basin 1 and Shelf 1 samples (Figure 2). MAG_00024 has six contigs and is dominated by reads from the Basin 2 and Shelf 2 samples (Figure 2). MAG_00029 is predicted to be 88.73% complete and 0% redundant, while MAG_00024 is 88.73% complete with 1.41% redundancy (Anvi’o stats). CheckM2 results further estimated 99.16% completeness and 0.09% contamination for MAG_00029 and 99.35% completeness and 0.08% contamination for MAG_00024 (Table S6).

**Figure 2.**
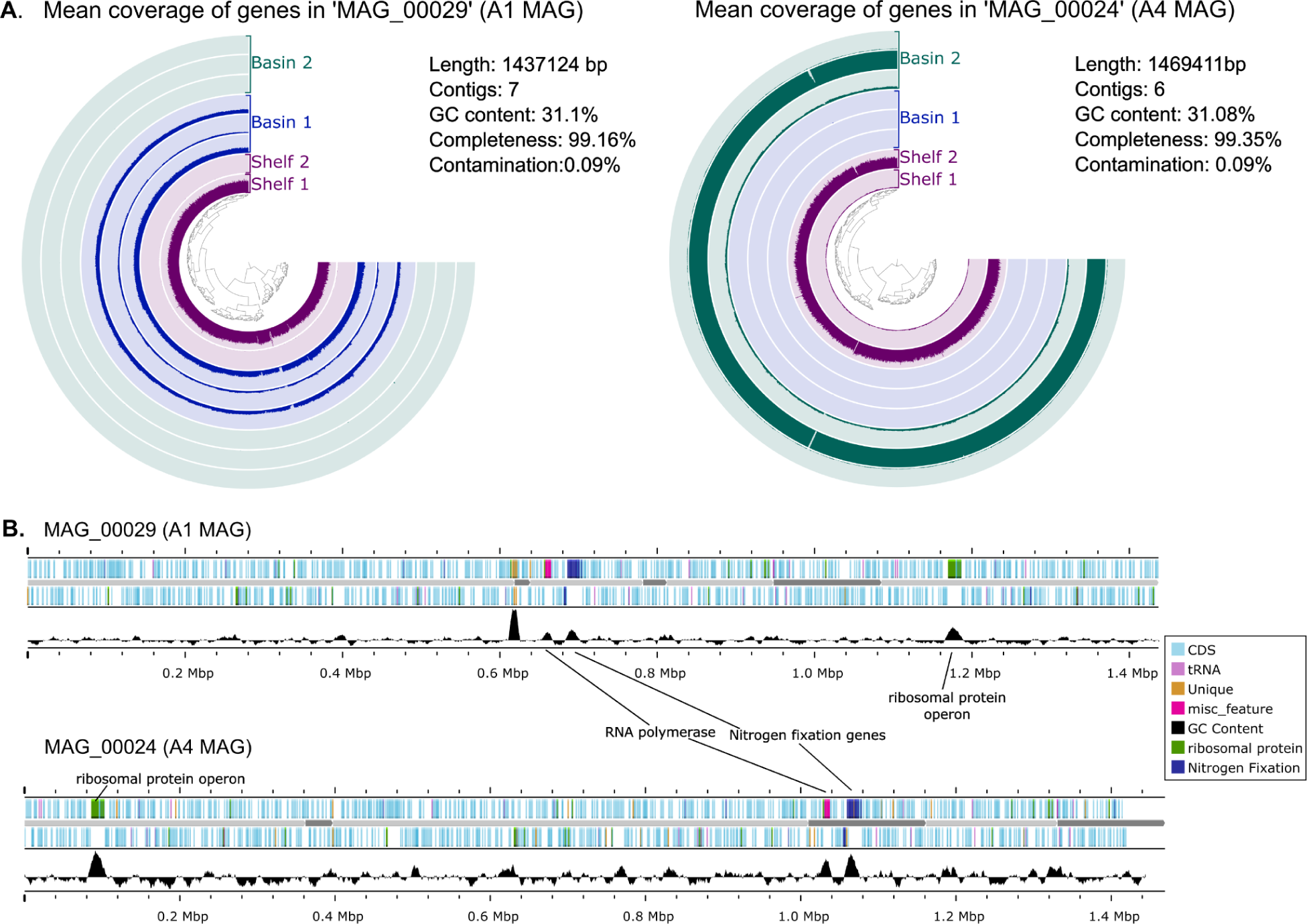
The two UCYN-A MAGs created from the metagenomic sequencing of enrichment culture water. (A) The average coverage of genes within each bin by sample shows MAG_00029 had a higher proportion of reads mapped from the Basin 1 and Shelf 1 samples. MAG_0024 showed a higher proportion of reads mapped from Basin 2 and Shelf 2 samples. (B) Annotations of MAG_00029 and MAG_00024 show areas of higher GC content aligning with the nitrogenase genes, two large RNA polymerase subunits, and a ribosomal protein operon.

*NifH* genes in both MAGs, aligned against other UCYN-A *nifH* ASVs and oligotypes, indicate MAG_00029 has 100% pairwise identity with the ASV ‘A1-deb’ (17) and the oligotype ‘Oligo_1’ (14), therefore this genome belongs to UCYN-A1. We will subsequently refer to MAG_00029 as the A1 MAG. The *nifH* gene from MAG_00024 has 100% pairwise identity with ASV ‘A4-511’ (17) and oligotype ‘Oligo_4’ (14), therefore MAG_00024 belongs to UCYN-A4 and will subsequently be referred to as the A4 MAG. Hence, our enrichment culturing and metagenomics was successful in obtaining A1 and A4 genomes from the coastal Northwest Atlantic.

### Pangenome comparisons and unique genes

The pangenome analysis revealed a high level of intra-sublineage conservation. The average nucleotide identity (ANI) was >99% between the three published UCYN-A1 genomes and between the four published UCYN-A2 genomes (Figure 3A; Table S7). In contrast, the ANI between A1 versus A2 was between 83.2-83.4%. The A4 MAG had 82.6-82.7% ANI versus the three A1 genomes and 85.2-85.3% ANI versus the four A2 genomes (Table S7), hence, the A4 genome is as different from each of A1 and A2 as these two are from each other.

**Figure 3.**
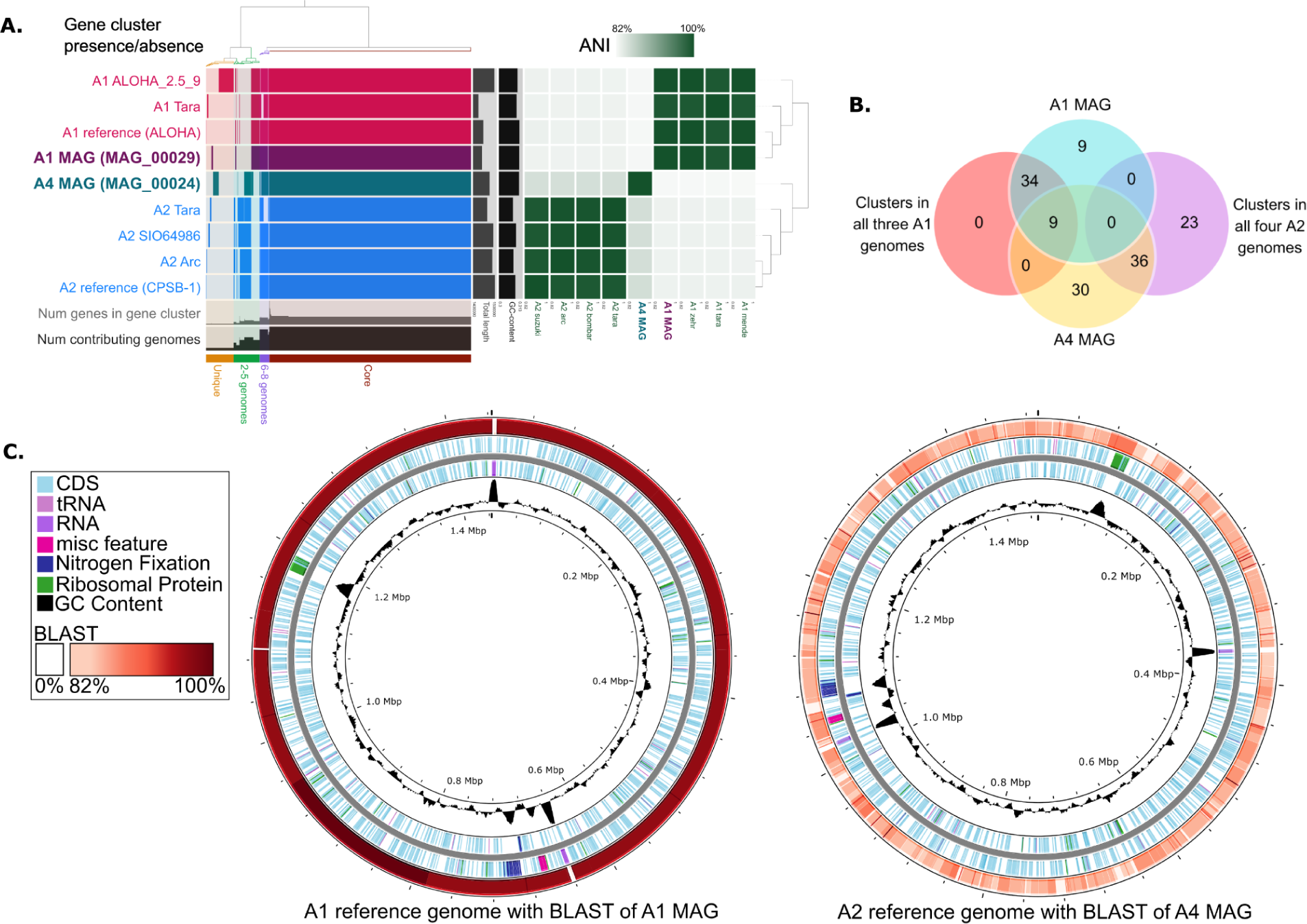
Pangenome analysis of UCYN-A. (A) Anvi’o schematic showing presence/absence of gene clusters in each genome (left) and average nucleotide identity (ANI) between each genome. (B) Venn-diagram of gene clusters in the MAGs that overlap with the clusters that are in all three published A1 genomes and all four published A2 genomes. (C) Full UCYN-A1 reference genome (A1 Zehr) annotated with BLAST results against MAG_00029 and full UCYN-A2 reference genome (A2 Suzuki) annotated with BLAST results against MAG_00024.

Analysis of gene cluster presence/absence resulted in 1,070 ‘core’ gene clusters [found in all nine genome], 52 ‘ambiguous’ gene clusters [in 6-8 genomes], 138 ‘other’ gene clusters [in 2-5 genomes], and 147 ‘unique’ gene clusters [in only one genome] (Figure 3A). Therefore, there are 337 peripheral gene clusters out of 1,407 total gene clusters. Of the 147 unique gene clusters, all but one contained a single gene (due to naming conventions in Anvi’o, note that in the analysis “clusters” can contain a single gene) (35). The A4 MAG has 30 unique gene clusters and genes, and the A1 MAG has 9 unique gene clusters and 10 unique genes (Figure 3B). Shared gene clusters between the published genomes include 43 and 59 shared exclusively between the three published UCYN-A1 genomes and the four A2 genomes, respectively. The A1 MAG possesses all of the 43 gene clusters present in all three of the published A1 genomes but none of those unique to the A2 genomes. The A4 MAG shares 9 of the 43 gene clusters unique to the three published A1 genomes and 36 of the 59 gene clusters unique to the four published A2 genomes (Figure 3B).

Of the 10 genes unique to the A1 MAG, five exhibit high sequence similarity to partial genes, hypothetical protein genes, or regions in the A1 reference genome with no genes (Table S8). Two of these unique genes had no annotation in Anvi’o and have no obvious counterparts using BLAST. The other three genes were annotated as a helicase conserved C-terminal domain coding region and two swr1 complex snf2 family DNA-dependent ATPases. These three show BLAST similarity with partial proteins from the genomes of the haptophytes *Emiliania huxleyi* and *Chrysochromulina tobinii* and were located in a high GC content (48%) area at the ends of two contigs with seven unique genes (Figure 2B). BLAST results of the A1 MAG against the A1 reference showed high percent identity of over 98% in most regions except for missing rRNA genes (Figure 3C).

Of the 30 gene clusters unique to the A4 MAG, only four were annotated by Anvi’o. Of the rest, three have BLAST matches to known proteins in public databases, eleven were annotated as hypothetical proteins, and twelve had no annotations beyond their original identification as ORFs (Table S9). Three of the genes with annotations are adjacent at the ends of two contigs: a *MalK* maltose/maltodextrin ATP-binding protein, a 23S rRNA-intervening sequence protein with BLASTx match to a four-helix bundle protein of unknown function from *Gracilimonas* sp., and a gene with a BLASTx match to a *HlyD* family efflux transporter from *Balneola* sp. The *MalK* has 93% identity in BLASTn with a portion of the same gene in the A2 reference genome, while the other two had no BLASTn results and do not align with the A2 reference genome. Other annotated genes encode an RNA polymerase sigma factor with BLASTx match to *Crocosphaera* sp., a glycosyltransferase *BscA*, a partial ferredoxin gene which partially aligns to the same gene in the A2 reference, and DUF3086 domain-containing protein of unknown function with a weak BLASTx match to the same protein in the A2 reference genome (45% amino acid identity). Five of the hypothetical protein genes and one unannotated gene were found to align with regions in the A2 reference genome with no annotations. A BLASTn of the A4 MAG against the A2 reference genome showed an average identity in the low 80% range and was also missing rRNA genes likely due to known difficulties in assembling the inverted repeated rDNA operon and associated genes (Figure 3C).

### Nitrogenase and 16S rRNA coding region comparison

Although the nif gene regions across sequenced genomes are largely syntenic (Figure 4), there are obvious differences that are likely due to a lack of complete genomes in some cases (e.g. a missing *nifH* gene in the A1 ALOHA_A2.5_9 genome (acc: GCA_022450625); see Supplementary Results S1 for specific details). Other differences unique to the A4 MAG are three hypothetical proteins that are not obviously present in any other genomes (Figure 4).

**Figure 4.**
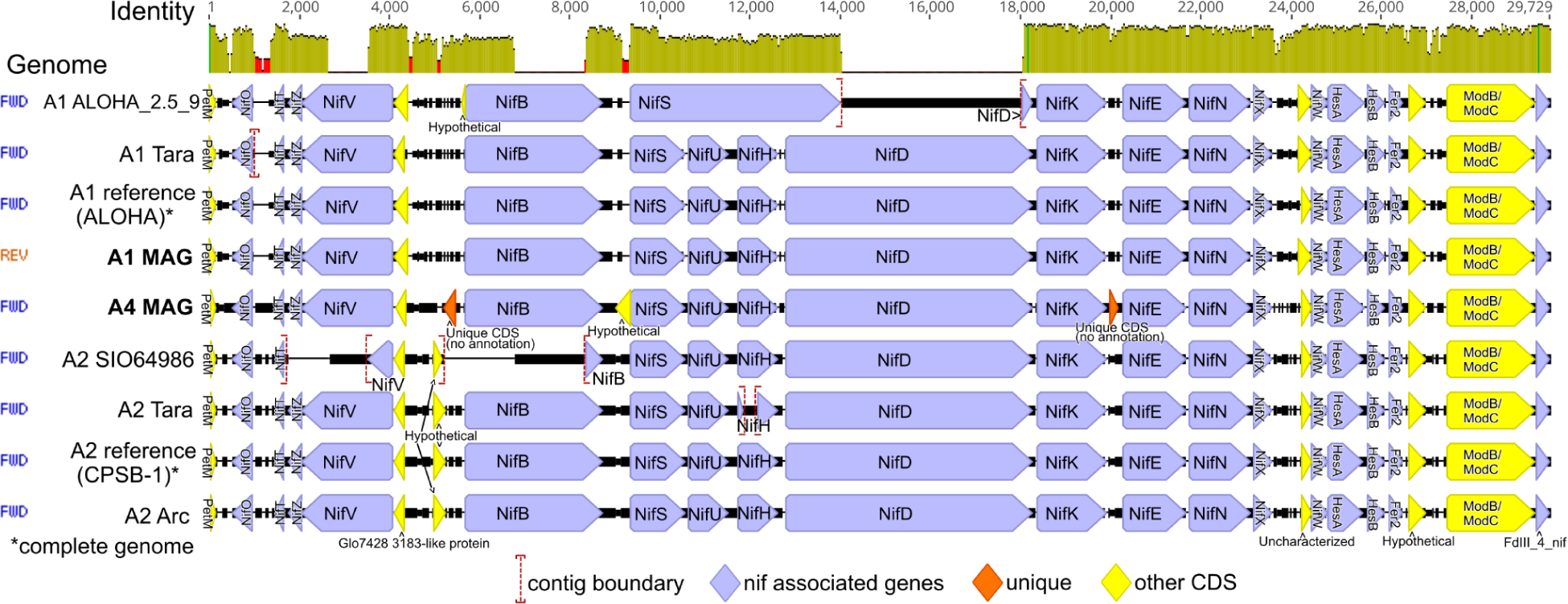
*Nif* gene region alignment of all genomes. includes the region between *PetM* and *FdIII_4_nif* genes chosen because they were present in all genomes. Genomes are ordered based on similarity and contig boundaries are marked in incomplete regions.

Two different 16S rRNA ASVs for UCYN-A were obtained by amplicon sequencing of our enrichment cultures: ASV022 (Basin 1 and Shelf 1) and ASV023 (Basin 2 and Shelf 2; Figure 1A-D). The genomes that contained the small subunit 16S rRNA gene were A1 the reference, A2 reference, A1 ALOHA_A2.5_9, A2 SIO64986, and the published A3 16S rRNA gene (MH807559) (38). ASV022 matched perfectly to the A1 reference and A1 ALOHA_A2.5_9 (Table S10). ASV023 was not 100% identical to any other 16S rRNA ASVs from published UCYN-A genomes; it was 99.2% identical to the 16S rRNA from the A2 SIO64986 and A3 genomes and 81.3% to the A2 reference sequence. The A2 reference 16S rRNA gene was the least similar to the other 16S genes in our analysis (Table S10).

### *B. bigelowii*/UCYN-A4 plastid genome

The *Braarudosphaera bigelowii* plastid genome assembled was only 1.5 kb smaller than the published chloroplast genome of *Chrysochromulina parva* (Figure 5A). Our new A4-associated plastid genome is incomplete since it lacks a fully assembled inverted repeat of the rRNA operon. Much of the published *C. parva* chloroplast genome is accounted for in homologous regions of our new *B. bigelowii* plastid genome, but there are many large-scale rearrangements, and some small unique regions exist in both plastid genomes (Figure 5B). Alignment analysis of the 16S rRNA derived from *B. bigelowii* plastids also suggests that intraspecific dissimilarities exist for this gene region within *B. bigelowii* (Supplementary Results S2; Table S11). BLAST results of aligning the *B. bigelowii* plastid genome (this study) with those recently published (OR912955.1, OR912954.1, OR912953.1) (13) showed 90-91% identity over partial regions of alignment (Figure S3).

**Figure 5.**
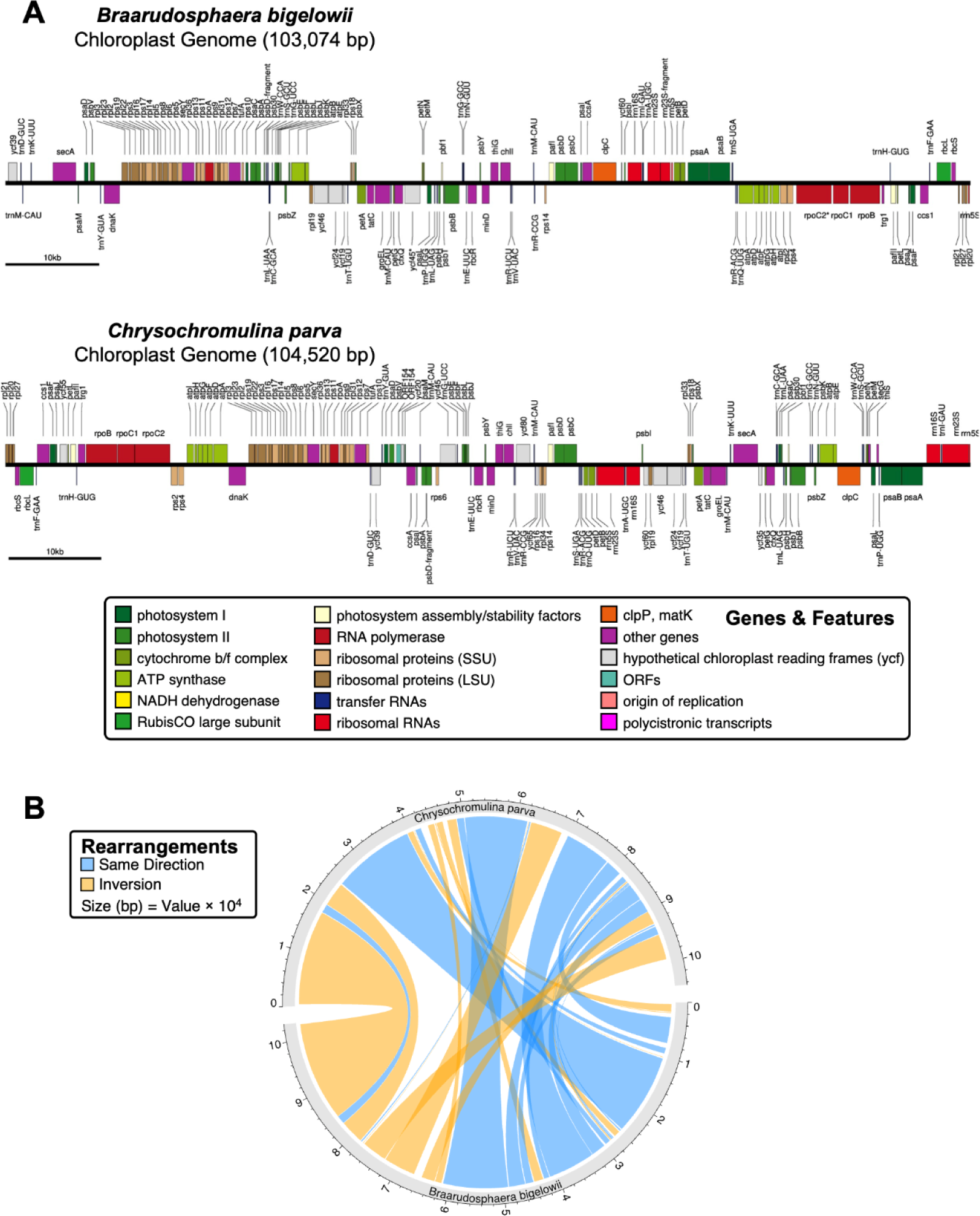
The *Braarudosphaera bigelowii* chloroplast genome annotated, shown above a previously published *Chrysochromulina parva* genome (A). The *B. bigelowii* genome has many of the same genes as the *C. parva* but is most noticeably missing the second rRNA operon which is difficult to assemble from metagenomic sequencing due to it being duplicated in the genome. Part B is a visualization of the MAUVE alignment of the two genomes which identifies homologous regions or ‘Locally Collinear Blocks’ and shows rearrangements.

## Discussion

### The A4 genome is more similar to the A2 genome and is hosted by a *B. bigelowii* genotype

The A4 MAG (MAG_00024) isolated from the coastal Northwest Atlantic is the first genome from the UCYN-A4 sublineage/ecotype reported, and the A1 MAG (MAG_00029) is the first one obtained from the Northwest Atlantic Ocean. The sequencing and assembly of MAGs is a powerful tool for exploring genomic diversity but can be problematic when closely related strains co-exist in samples (50). Pangenome results suggest that this is not the case for the UCYN-A MAGs presented herein and give evidence that they are genomes of separate sublineages. Although UCYN-A2 is known to be prevalent in the coastal NW Atlantic and can co-occur with A4 (17), we did not detect it in the enrichment cultures, nor did we detect the A2 host *B. bigelowii* ASV. The A4 MAG shows more similarities to the A2 ecotype than the A1 ecotype; it has slightly higher ANIs with the A2 genomes and overlaps with more sublineage-specific gene clusters from A2 (Figure 3A, B). All analysis conducted in this study including 16S rRNA metabarcoding, *nif* comparisons, and pangenomics support the phylogenetic relationships which show that UCYN-A4 is more closely related to A2 than to A1 (Figure S1). This suggests that UCYN-A4 diverged from A2 after A1 and A2 diverged. With fewer host sequences, it is harder to see a pattern of evolution, but if the symbiotic partnership between the two organisms is obligate, one would expect the evolution of the host to mirror that of UCYN-A. The ANI comparison, number of unique genes, and the *nif* gene coding regions (Figure 3, 4) together also support the A4 MAG as a distinct sublineage because its genome is sufficiently different from both A1 and A2.

The presence of a corresponding *B. bigelowii* 16S rRNA ASV, 18S rRNA ASV, and near-complete plastid genome in enrichment cultures alongside an A4 genome supports the hypothesis that a *B. bigelowii* algae of Genotype I hosts the A4 ecotype and these genetic signatures are likely evolutionary pairs. Meanwhile, our samples with UCYN-A1 contained a different *B. bigelowii* 16S rRNA [ASV020] sequence, pointing also to a *B. bigelowii* genotype as the possible host signature for the A1 MAG identified herein different from the A2 and A4 hosts. Intriguingly, this 16S rRNA ASV is different from the *Chrysochromulina* sp. ASV found commonly co-occurring in the Bedford Basin with UCYN-A1 (17). Studies have shown that *B. bigelowii* has been observed in culture without UCYN-A (37), so it is possible that the prior *Chrysochromulina* sp. was not harboring UCYN-A1 but was simply co-occurring.

### Genomes of UCYN-A in the Pacific and Atlantic Oceans share high identity

The A1 MAG had an identical full *nifH* gene to the A1 reference, and 99.9% ANI, making the two effectively indistinguishable even though they are from different ocean basins (Pacific versus Atlantic); these results support findings of previous sublineage comparisons (51). Unique A1 MAG genes occur at the ends of contigs and therefore may be erroneous due to lower coverage or repeats. It is likely that these occur where the rRNA operons are missing. RNA genes are difficult to assemble and bin into MAGs since they are highly conserved, have different GC content, and often have multiple copies in a genome, such as in UCYN-A where there are two rRNA operons (52). At this unique region in the A1 MAG, there is higher GC content, and it is located adjacent to the N_2_ fixation gene region and RNA polymerase genes, similar to the location of rRNA genes present in the A1 reference genome (Figure 2B) (36). Therefore, it can be hypothesized that this region is just missing the rRNA operon leading to larger differences from the reference genome.

### Co-enriched microbial groups

Microbes present at high percent relative abundances in the enrichment cultures largely represent those occurring when *B. bigelowii* and UCYN-A are present in their natural habitat as well (Figure 1E). Though the ASVs were not exactly the same between cultured samples, the similarity of co-enriched organisms at higher taxonomic levels (e.g., genera) gives some initial insight into which microbial groups might occur together with *B. bigelowii*/UCYN-A. For example, some haptophytes are known to be mixotrophic (53,54) and prior studies support the idea that *B. bigelowii* is capable of phagocytosis both as the mechanism for it acquiring UCYN-A originally as an endosymbiont and for acquiring nutrients (37). Although still untested, it is conceivable that some of the microbes that co-enriched with UCYN-A may be a food source for mixotrophic *B. bigelowii* (55), which may have in-turn helped this haptophyte remain in our cultures for multiple weeks. Alternatively, these microbes may have been enriched alongside *B. bigelowii* simply because their growth was stimulated by the same temperature and lighting conditions and/or they may have been stimulated via the use of carbon and nitrogen that would have theoretically been fixed by *B. bigelowii*/UCYN-A present in the cultures (note we did not add nitrogen or carbon nutrients to any of our incubated seawater (56)). Given that some of the same bacteria were retained after targeting larger eukaryotic cells that contained UCYN-A *nifH* sequences during FACS, these microbes may have also been physically co-sorted as single events/particles (e.g., *Pelagibacter ubique*; Figure 1B). In general, now that *B. bigelowii*/UCYN-A2 has been cultured (13) future studies could be aimed at better resolving more specific interspecies dynamics and metabolic interactions between *B. bigelowii* and other marine microbes. However, given the vast amount of microbial diversity present in the ocean (57), having a narrower list of microbes known to co-culture *ex situ* and/or co-occur in the natural environment with *B. bigelowii–*like those mentioned herein and also identified from prior network analyses (17,58,59)–can help inform the selection of microbial strains for this type of future research.

### UCYN-A4 as a possible nitroplast

UCYN-A2 was classified as a nitroplast because it has the combination of the necessary characteristics: a reduced genome, cell architecture integration and synchronous division with other organelles, and imported proteins from the host (13). The A4 MAG has the same genome reduction that is characteristic of UCYN-A with a length within the same 1.4 Mbp range. It is near complete but is missing the two rRNA operons likely for the same reasons as the A1 MAG above. Annotated genes unique to the A4 MAG were mostly hypothetical genes that align only partially to the A2 reference genome, or located on the ends of contigs which are error prone regions due to the assembly process (Table S9). Protein import is considered to be a defining feature of an organelle relative to an endosymbiont (60). Such proteins imported into the nitroplast from the host complemented key steps in biological pathways that are missing genes in the A2 genome because of genome reduction (13). We found that all of these missing genes in A2 are also missing in the previously published A1 reference genome (40) and from the A4 MAG. This supports the idea that host-encoded proteins may be filling in these gaps in the other sublineages as well. It is important to note that the genomes of UCYN-A sublineages and their associated algal chloroplast genes are different enough to warrant separate investigation of each to determine their place on the evolutionary path to becoming organelles.

Relative to the endosymbiotic origins of mitochondria and plastids from alphaproteobacteria and cyanobacteria respectively (61), the UCYN-A2 nitroplast of *B. bigelowii* appears to be at a much earlier stage of organellogenesis, similar to that seen in the chromatophore of the testate amoeba *Paulinella* (13,62). Phylogenomic studies suggest that the common UCYN-A ancestor was already associated with an ancestor of *B. bigelowii*, and had already undergone genome reduction, before the A1 and A2 lineages diverged ∼90 Myr ago (63,64). It is unclear at what stage of organellogenesis UCYN-A was at the time of divergence, and unclear now as to whether the sublineages of UCYN-A are all at the same stage. With further study, the comparison of the plastid genomes between hosts can be done in parallel with the comparison of UCYN-A/nitroplast genomes and opens the door for studying the details of their evolution and divergence. Using new culturing techniques plus long-read sequencing, the full nuclear and organellar genomes of different genotypes can be compared to determine the details of each UCYN-A-containing *B. bigelowii* lineage.

## Acknowledgments

We would like to thank the Bedford Institute of Oceanography for welcoming our weekly participation in the Bedford Basin Monitoring Program as well as all of those who have helped with Bedford Basin sampling including the Ocean Frontier Institute technical team. We would also like to thank those who sampled during the 2021 MORI Atlantic Condor Expedition and the Integrated Microbiome Resource and specifically André M. Comeau for providing feedback with respect to sequencing strategy. E.J.H.K. was funded through the Ocean Frontier Institute’s Ocean Graduate Excellence Network. Funding for J.L.R was provided from the Canada First Research Excellence Fund, Ocean Frontier Institute Module C, and a Discovery Grant. Support from the Archibald Lab was provided by an Arthur B. McDonald Chair and a Discovery Grant (RGPIN-2019-05058). During experiments B.M.R. was supported by the Natural Sciences and Engineering Research Council of Canada (NSERC) through a Canada Graduate Scholarship-Doctoral Award; B.M.R. is also currently supported by NSERC via a Postdoctoral Fellowship. A Graduate Scholarship and a Killam Predoctoral Award from Dalhousie University further supported B.M.R. during the study.

## Data availability statement

Data is available upon request

