## Supplementary text and tables for "Targeted metagenomics reveals pangenomic diversity of the nitroplast (UCYN-A) and its algal host plastid"

**Supplementary Information**

Supplementary Text:

- Supplementary Methods S1. Commands used to create contigs database
- Supplementary Methods S2. Commands used for Pangenome analysis
- Supplementary Methods S3. Plastid Annotation and Synteny
- Supplementary Methods S3. Plastid Annotation and Synteny
- Supplementary Results S1. Specific details on *nifH* gene region differences
- Supplementary Results S2. 16S rRNA comparisons from *B. bigelowii* plastids

Supplementary Tables:

- Table S1. Information about all genomes in pangenome analysis
- Table S2. Table of UCYN-A enrichment culturing sample methods details.
- Table S3. Metagenomic sequencing details for each enrichment culturing sample.
- Table S4. Table of the top 0.1% of all 16S ASVs used in figures, names, taxonomy, DNA sequence (tableS4_16S_topASVs.xlsx).
- Table S5. Table S5. (A) 18S rRNA data used in alignment. Accessions, genotype, sources. (B) Distance matrix of identities of alignment (tableS5_18S.xlsx).
- Table S6. CheckM2 Results of genomes in the pangenome analysis
- Table S7. Matrix of ANI values between each genome in the pangenome
- Table S8. Table of unique genes to the A1 MAG (tableS8_S9_uniqueGenes.xlsx).
- Table S9. Table of unique genes to A4 MAG (tableS8_S9_uniqueGenes.xlsx).
- Table S10. Distance matrix of UCYN-A16S rRNA gene sequence identities from alignment.
- Table S11. Distance matrix of plastid 16S rRNA gene sequence identities from alignment (*this study).

Supplementary Figures:

- Figure S1. Full tree of all 186 ASVs present in all four enrichment cultures above 0.1% relative abundance.
- Figure S2. Alignment of NifH amino acid sequences from each of the genomes where *nifH* gene was present.
- Figure S3. Alignment of our *B. bigelowii* plastid sequence with published *B. bigelowii* chloroplast sequence from Coale et al. (13) (acc: OR912955.1, OR912954.1, OR912953.1).

**Supplementary Methods S1. Commands used to create contigs database:**
Prior to the first step in the pipeline, quality filtering of raw reads was done using the 'iu-filter-quality-minoche' program from the illumina-utils library v1.4.1 (65). Co-assembly was done using MEGAHIT v1.0.3 (66) and mapping of short reads onto contigs was done using Bowtie2 v2.0.5 (67). Contig fasta files were then reformatted and 'anvi-gen-contigs-database' was further run to generate a contigs database including open reading frames identified by Prodigal v2.6.3 (25,68).

**Supplementary Methods S2. Commands used for Pangenome analysis:**
Using the contigs database created specifically for the pangenome analysis, a 'genomes storage database' was generated from all the genomes and analysis was run using the command 'anvi-pan-genome' with parameters '--use-ncbi-blast', '--minbit 0.5', and '--mcl-inflation 10'. This program (35) calculates the similarities between genomes, identifies and counts gene clusters, and stores this information in a pan-database to be used for analysis and visualization. Calculation of average nucleotide identity (ANI) was done using the 'anvi-compute-genome-similarity' command with the program PyANI v0.2 (69). Pangenome visualizations were created using the 'anvi-display-pan' command (35).

**Supplementary Methods S3. Plastid Annotation and Synteny:**
Both the B. bigelowii UCYN-A4 containing plastid genome and C. parva genomes were annotated using GeSeq with the tools: ARAGORN v1.2.38, blatN, and blatX (70–72). To assess synteny, the B. bigelowii plastid genome was further aligned to that of C. parva using MAUVE v1.1.3 (46) and visualized
in R (v4.3.0) via RStudio v2023.06.1+524 using the circlize package v0.4.16 (73).

**Supplementary Methods S3. Plastid Annotation and Synteny**:
16S rRNA amplicon sequencing results were transformed into percent abundances and plotted in R using packages tidyverse v2.0.0 and ggplot2 v3.5.0 (74). Taxonomic assignment of amplicon sequence variants (ASVs) with >0.1% abundance in each culture was manually refined following the same methods as Robicheau et al. (23). 16S rRNA amplicon sequence data from the Bedford Basin weekly time series was from the same original dataset and methods as in Robicheau et al. (23) with the exception that all ASVs were retained during analysis rather than subsetting for only plastid signatures. ASVs with >1% relative abundance in the four enrichment cultures were extracted from the time series dataset and plotted to show their annual weekly relative abundance in the natural seawater of the Bedford Basin. A neighbor-joining tree with 1000 bootstraps for 16S rRNA ASVs in the enrichment cultures was built using the online PhyML tool (75) and Muscle v3.8.625 to align sequences (76).

**Supplementary Results S1. Specific details on NifH gene region differences:**
The A2 SIO64986 and A1 ALOHA_A2.5_9 genomes both lack a complete suite of nitrogen fixation genes. The A2 SIO64986 genome does have nifH, while in the A1 ALOHA_A2.5_9 genome an obvious nifH gene is not present (Figure 4). The nifH from the A2 Tara genome is split between two contigs and is missing the portion which aligns to known ASVs or oligotypes. All nifH genes found in the A1 and A2 genomes have 100% pairwise identity to the sequences 'A1-deb' and 'Oligo_1 and 'A2-bdd' and 'Oligo_3' (14,17), respectively.
Comparison of the nifH-derived amino acid sequences from each genome shows twelve positions with disagreements mainly in the last 10 amino acids (Figure S2). The A1 reference genome's NifH amino acid sequence was identical to that of the A1 MAG (Figure S2).
Alignment of the full Nif gene region showed a large syntenic block containing 23 genes (Figure 4). There was also one hypothetical protein in the same location for the three published A2 genomes. As mentioned, the A4 MAG has three hypothetical proteins that are not obviously present in any other genomes. For the A2 SIO64986 and A1 ALOHA_A2.5_9 genomes this region spans multiple contigs, so there are fragmented gene regions thus making it harder to align these accurately.

**Supplementary Results S2. 16S rRNA comparisons from B. bigelowii plastids:**

The 16S rRNA gene from our B. bigelowii plastid genome was aligned to other published B. bigelowii plastid 16S rRNA genes sequences, as well as to those from other haptophyte plastids and 16S rRNA ASVs obtained herein. All of the B. bigelowii 16S rRNA genes analyzed are >99% identical to each other (Table S11). Our B. bigelowii plastid 16S rRNA gene was 100% identical to the B. bigelowii ASV021, which was in the Shelf 2 and Basin 2 amplicon sequencing results. The ASV020 which was in Shelf 1 and Basin 1 was not 100% identical to any of the others, but it had >99% identity to the other B. bigelowii 16S rRNA plastid sequences (Table S11).

**Supplementary Tables:**

| Genome Name | Sublineage | Location | Number of Contigs | Length (bp) | Assembly Accession | File Name | Reference |
|---|---|---|---|---|---|---|---|
| A1 reference (ALOHA) | A1 | 22.75 N, 158 W | Complete Genome | 1443806 | GCA_000025125.1 | | Tripp et al. (36) |
| A1 ALOHA_A2.5_9 | A1 | 22.75 N, 158 W | 47 | 1489669 | GCA_022450625.1 | | Leu et al. (77) |
| A1 Tara | A1 | 21.4788 S, 56.8291 E | 44 | 1422642 | | TARA_IOS_50_MAG_00026 | Delmont et al. (78) |
| A2 reference (CPSB-1) | A2 | 35.5171 N, 133.9374 E | Complete Genome | 1491611 | GCA_020885515.1 | | Suzuki et al. (37) |
| A2 SIO64986 | A2 | 32.87 N, 117.25 W | 52 | 1485499 | GCA_000737945.1 | | Bombar et al. (63) |
| A2 Arc | A2 | 67.1705 N, 0.4423 E | 5 | 1480855 | | Arc-UCYN-A2 | Shiozaki et al. (51) |
| A2 Tara | A2 | 47.1863 S, 58.2902 W | 46 | 1459650 | | TARA_AOS_82_MAG_00023 | Delmont et al. (78) |
| A1 MAG | A1 | 42.50 N, 61.43 W | 7 | 1437124 | | RUN1_ALL_MAG_00029 | this study |
| A4 MAG | A4 | 42.50 N, 61.43 W | 6 | 1469411 | | RUN1_ALL_MAG_00024 | this study |

Table S1. Information about all genomes in pangenome analysis

| Sample | | Date | Initial Incubation Nutrients | Sorted Cells | Location | Coordinates | Depth | qPCR | Total 16S V6V8 |
|---|---|---|---|---|---|---|---|---|---|
| Basin 1 | 2018_BB_a<br>2018_BB_b<br>2018_BB_c | 2018 | 2nM Fe + 200nM PO4, 13 weeks at 15℃ with 12h/12h light/dark cycle | No; Seawater incubated with Nutrients straight to sequencing post DNA-extraction | Bedford Basin | 44º 41' 37" N, 63º 38' 25" W | Surface | Non-specific SYBR qPCR Assay; Had UCYN-A | Similar to A1 |
| Basin 2 | 2020_BB_a<br>2020_BB_b<br>2020_BB_c | 2020 | 2nM Fe + 400 nM PO4 + 0.5ml/L vitamins, 2 weeks at 15℃ with 12h/12h light/dark cycle | Cells Sorted (2,000 cells) + Incubation for 8.5 weeks in ~10mL of 0.2um filtered seawater from original sample. Enriched with more nutrients at week 0,2, and 4. | Bedford Basin | 44º 41' 37" N, 63º 38' 25" W | 5m | Used A1 and A2 TaqMan Assays; Mainly A2 | Similar to A2 |
| Shelf 1 | 2021_Shelf_a | 2021 | 2nM Fe + 200nM PO4 | No; Seawater incubated with Nutrients straight to sequencing post DNA-extraction | Scotian Shelf | 42º 50' 00" N, 61º 43' 00" W | 20m | Ecotype Specific TaqMan Assays; ~7.5:1 for A1:A2 | UCYN-A ASV with higher % similar to A1 |
| Shelf 2 | 2021_Shelf_b | 2021 | 2nM Fe + 200nM PO4 | Cells sorted without further Incubation (28,911 cells) | Scotian Shelf | 42º 50' 00" N, 61º 43' 00" W | 20m | Ecotype Specific TaqMan Assays; A2>>A1 | Similar to A2 |

Table S2. Table of UCYN-A enrichment culturing sample methods details.

| Additional sequencing info | Technology - Run | Depth | Date | R1 | R2 |
|---|---|---|---|---|---|
| 2018_BB_a | MiSeq - MS251 (S289) | 1X | 5/3/2019 | BBCulture9-20180822_S289_L001_R1_001.fastq.gz | BBCulture9-20180822_S289_L001_R2_001.fastq.gz |
| 2018_BB_b | NextSeq - NS39 | 4X | 5/30/2019 | Robichaud_S88_L001_R1_001.fastq.gz, Robichaud_S88_L002_R1_001.fastq.gz, Robichaud_S88_L003_R1_001.fastq.gz, Robichaud_S88_L004_R1_001.fastq.gz | Robichaud_S88_L001_R2_001.fastq.gz, Robichaud_S88_L002_R2_001.fastq.gz, Robichaud_S88_L003_R2_001.fastq.gz, Robichaud_S88_L004_R2_001.fastq.gz |
| 2018_BB_c | MiSeq - MS212 (S385) | 1X | 12/21/2018 | BBculture9_S385_L001_R1_001.fastq.gz | BBculture9_S385_L001_R2_001.fastq.gz |
| 2020_BB_a | NextSeq - NS79 (S42) | 4X | 2/7/2022 | BBc11-20200916-5m-UA2g_S42_L001_R1_001.fastq.gz, BBc11-20200916-5m-UA2g_S42_L002_R1_001.fastq.gz, BBc11-20200916-5m-UA2g_S42_L003_R1_001.fastq.gz, BBc11-20200916-5m-UA2g_S42_L004_R1_001.fastq.gz | BBc11-20200916-5m-UA2g_S42_L001_R2_001.fastq.gz, BBc11-20200916-5m-UA2g_S42_L002_R2_001.fastq.gz, BBc11-20200916-5m-UA2g_S42_L003_R2_001.fastq.gz, BBc11-20200916-5m-UA2g_S42_L004_R2_001.fastq.gz |
| 2020_BB_b | NextSeq - NS83 | 4X | 3/16/2022 | BBc11-20200916-5m-UA2g_S46_L001_R1_001.fastq.gz, BBc11-20200916-5m-UA2g_S46_L002_R1_001.fastq.gz, BBc11-20200916-5m-UA2g_S46_L003_R1_001.fastq.gz, BBc11-20200916-5m-UA2g_S46_L004_R1_001.fastq.gz | BBc11-20200916-5m-UA2g_S46_L001_R2_001.fastq.gz, BBc11-20200916-5m-UA2g_S46_L002_R2_001.fastq.gz, BBc11-20200916-5m-UA2g_S46_L003_R2_001.fastq.gz, BBc11-20200916-5m-UA2g_S46_L004_R2_001.fastq.gz |
| 2020_BB_c | NextSeq - NS72 (S69) | 1X | 8/27/2021 | BBc11_20200916_5m_UA2g_S69_L001_R1_001.fastq.gz, BBc11_20200916_5m_UA2g_S69_L002_R1_001.fastq.gz, BBc11_20200916_5m_UA2g_S69_L003_R1_001.fastq.gz, BBc11_20200916_5m_UA2g_S69_L004_R1_001.fastq.gz | BBc11_20200916_5m_UA2g_S69_L001_R2_001.fastq.gz, BBc11_20200916_5m_UA2g_S69_L002_R2_001.fastq.gz, BBc11_20200916_5m_UA2g_S69_L003_R2_001.fastq.gz, BBc11_20200916_5m_UA2g_S69_L004_R2_001.fastq.gz |
| 2021_Shelf_a | NextSeq - NS99 | 6X | 12/6/2022 | BR2021_HL7_20m_enriched_S86_L001_R1_001.fastq.gz | BR2021_HL7_20m_enriched_S86_L001_R2_001.fastq.gz |
| 2021_Shelf_b | NextSeq - NS99 | 6X | 12/6/2022 | JA_SortedCells_UA2_rep10_S87_L001_R1_001.fastq.gz | JA_SortedCells_UA2_rep10_S87_L001_R2_001.fastq.gz |

Table S3. Metagenomic sequencing details for each enrichment culturing sample.

Table S4. Table of the top 0.1% of all 16S ASVs used in figures, names, taxonomy, DNA sequence (tableS4_16S_topASVs.xlsx).

Table S5. (A) 18S rRNA data used in alignment. Accessions, genotype, sources. (B) Distance matrix of identities of alignment (tableS5_18S.xlsx).

| Genome Name | Completeness | Contamination | Completeness Model Used | Translation Table Used | Coding Density | Contig N50 | Average Gene Length | Genome Size | GC Content | Total Coding Sequences | Additional Notes |
|---|---|---|---|---|---|---|---|---|---|---|---|
| A1 ALOHA_2.5_9 | 99.22 | 0.52 | Neural Network (Specific Model) | 11 | 0.817 | 86550 | 316.285047 | 1489669 | 0.31 | 1284 | None |
| A1 Tara | 98.2 | 0.19 | Neural Network (Specific Model) | 11 | 0.816 | 51922 | 320.384934 | 1422642 | 0.31 | 1208 | None |
| A1 reference (ALOHA) | 99.58 | 0.09 | Neural Network (Specific Model) | 11 | 0.808 | 1443806 | 325.370401 | 1443806 | 0.31 | 1196 | None |
| A2 Arc | 99.17 | 0.1 | Neural Network (Specific Model) | 11 | 0.791 | 364024 | 321.712521 | 1480855 | 0.31 | 1214 | None |
| A2 SIO64986 | 98.62 | 0.09 | Neural Network (Specific Model) | 11 | 0.785 | 73988 | 311.590545 | 1485499 | 0.31 | 1248 | None |
| A2 reference (CPSB-1) | 99.45 | 0.09 | Neural Network (Specific Model) | 11 | 0.785 | 1491611 | 322.82562 | 1491611 | 0.31 | 1210 | None |
| A2 Tara | 96.47 | 0.38 | Neural Network (Specific Model) | 11 | 0.788 | 44130 | 316.413366 | 1459650 | 0.31 | 1212 | None |
| A4 MAG | 99.35 | 0.08 | Neural Network (Specific Model) | 11 | 0.802 | 361971 | 325.426325 | 1469411 | 0.31 | 1208 | None |
| A1 MAG | 99.16 | 0.09 | Neural Network (Specific Model) | 11 | 0.817 | 352113 | 327.367057 | 1437124 | 0.31 | 1196 | None |

Table S6. CheckM2 results of genomes in the pangenome analysis

| Genome Name | A1 reference (ALOHA) | A1 ALOHA_A2.5_9 | A1 Tara | A2 reference (CPSB-1) | A2 SIO64986 | A2 Arc | A2 Tara | A1 MAG | A4 MAG |
|---|---|---|---|---|---|---|---|---|---|
| **A1 ALOHA_A2.5_9** | 100.00 | 99.25 | 99.93 | 83.26 | 83.34 | 83.37 | 83.24 | 99.90 | 82.76 |
| **A1 Tara** | 99.23 | 100.00 | 99.26 | 83.30 | 83.30 | 83.34 | 83.29 | 99.28 | 82.74 |
| **A1 reference (ALOHA)** | 99.92 | 99.28 | 100.00 | 83.32 | 83.44 | 83.46 | 83.28 | 99.91 | 82.72 |
| **A2 Arc** | 83.27 | 83.32 | 83.33 | 100.00 | 99.63 | 99.33 | 99.74 | 83.33 | 85.34 |
| **A2 SIO64986** | 83.37 | 83.31 | 83.42 | 99.62 | 100.00 | 99.26 | 99.65 | 83.31 | 85.33 |
| **A2 reference (CPSB-1)** | 83.38 | 83.34 | 83.42 | 99.33 | 99.25 | 100.00 | 99.31 | 83.32 | 85.40 |
| **A2 Tara** | 83.31 | 83.35 | 83.36 | 99.73 | 99.64 | 99.30 | 100.00 | 83.36 | 85.35 |
| **A4 MAG** | 99.90 | 99.30 | 99.91 | 83.27 | 83.26 | 83.28 | 83.28 | 100.00 | 82.75 |
| **A1 MAG** | 82.60 | 82.69 | 82.65 | 85.26 | 85.31 | 85.28 | 85.33 | 82.65 | 100.00 |

Table S7. Matrix of ANI values between each genome in the pangenome analysis.

Table S8. Unique genes to the A1 MAG (tableS8_S9_uniqueGenes.xlsx).

Table S9. Unique genes to the A4 MAG (tableS8_S9_uniqueGenes.xlsx).

| 16S Sequence | A2 reference - 16S rRNA SSU | A1 ALOHA_A2.5_9 - 16S rRNA SSU | A1 reference - 16S rRNA SSU | Basin 1 ASV022 | A3 - partial 16S rRNA | A2 SIO64986 - 16S rRNA SSU | Shelf 2 ASV023 |
|---|---|---|---|---|---|---|---|
| A2 reference - 16S rRNA SSU | | 88.072 | 88.067 | 80.789 | 86.825 | 88.27 | 81.316 |
| A1 ALOHA_A2.5_9 - 16S rRNA SSU | 88.072 | | 100 | 100 | 98.946 | 98.662 | 97.368 |
| A1 reference - 16S rRNA SSU | 88.067 | 100 | | 100 | 98.946 | 98.714 | 97.368 |
| Basin 1 ASV022 | 80.789 | 100 | 100 | | 97.632 | 98.158 | 97.368 |
| A3 - partial 16S rRNA | 86.825 | 98.946 | 98.946 | 97.632 | | 98.645 | 99.211 |
| A2 SIO64986 - 16S rRNA SSU | 88.27 | 98.662 | 98.714 | 98.158 | 98.645 | | 99.211 |
| Shelf 2 ASV023 | 81.316 | 97.368 | 97.368 | 97.368 | 99.211 | 99.211 | |

Table S10. Distance matrix of UCYN-A16S rRNA gene sequence identities from alignment.

| Plastid 16S rRNA sequence | KJ201907 – C. tobinii 16S rRNA | NC_036937 – C. parva 16S rRNA | AY741371 – Ehux 16S rRNA | LC595682 – NIES-4442 | BRA4 – B. bigelowii - 16S rRNA* | 16S ASV021 – B. bigelowii | AB847984 – isolate TMRscBb1 | AB847985 – isolate TMRscBb7 | AB847986 – isolate TMRscBb8 | 16S ASV020 – B. bigelowii |
|---|---|---|---|---|---|---|---|---|---|---|
| KJ201907 – C. tobinii 16S rRNA | | 100 | 94.494 | 94.868 | 94.969 | 91.579 | 94.873 | 94.895 | 94.873 | 91.316 |
| NC_036937 – C. parva 16S rRNA | 100 | | 94.494 | 94.868 | 94.963 | 91.579 | 94.873 | 94.895 | 94.873 | 91.316 |
| AY741371 – Ehux 16S rRNA | 94.494 | 94.494 | | 95.423 | 95.649 | 92.368 | 95.745 | 95.694 | 95.673 | 92.632 |
| LC595682 – NIES-4442 | 94.868 | 94.868 | 95.423 | | 99.584 | 99.474 | 99.963 | 99.963 | 99.963 | 99.737 |
| BRA4 – B. bigelowii - 16S rRNA* | 94.969 | 94.963 | 95.649 | 99.584 | | 100 | 99.673 | 99.691 | 99.673 | 99.737 |
| 16S ASV021 – B. bigelowii | 91.579 | 91.579 | 92.368 | 99.474 | 100 | | 99.474 | 99.474 | 99.474 | 99.737 |
| AB847984 – isolate TMRscBb1 | 94.873 | 94.873 | 95.745 | 99.963 | 99.673 | 99.474 | | 99.909 | 99.964 | 99.737 |
| AB847985 – isolate TMRscBb7 | 94.895 | 94.895 | 95.694 | 99.963 | 99.691 | 99.474 | 99.909 | | 99.909 | 99.737 |
| AB847986 – isolate TMRscBb8 | 94.873 | 94.873 | 95.673 | 99.963 | 99.673 | 99.474 | 99.964 | 99.909 | | 99.737 |
| 16S ASV020 – B. bigelowii | 91.316 | 91.316 | 92.632 | 99.737 | 99.737 | 99.737 | 99.737 | 99.737 | 99.737 | |

Table S11. Distance matrix of plastid 16S rRNA gene sequence identities from alignment (*this study).
