## Supplemental Figures for "Targeted metagenomics reveals pangenomic diversity of the nitroplast (UCYN-A) and its algal host plastid"

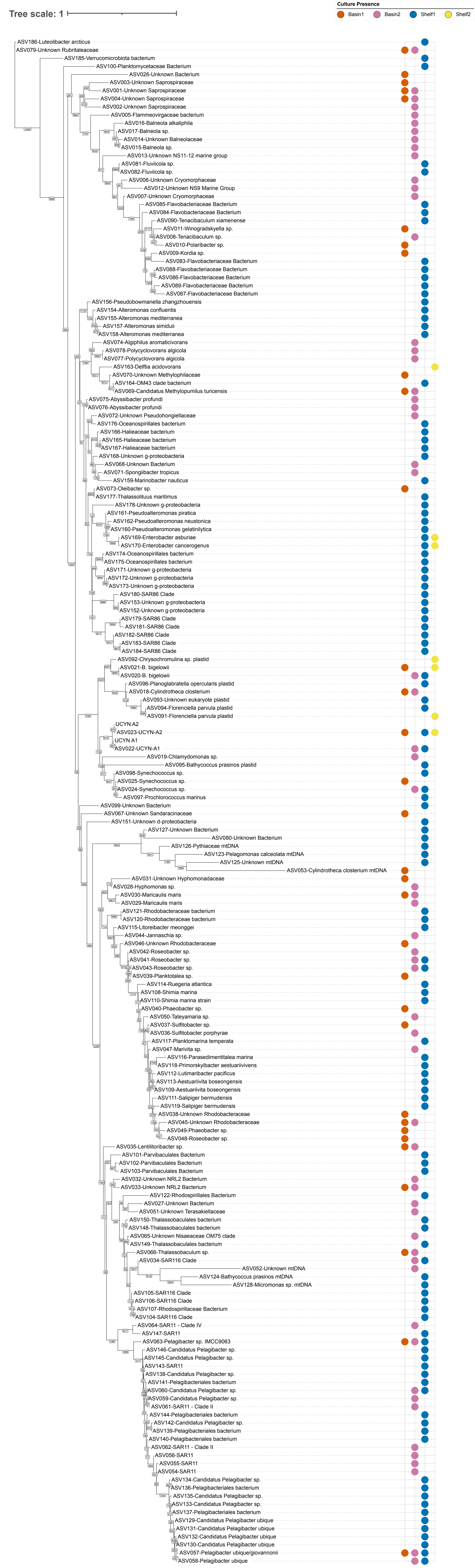


Figure S1. Full tree of all ASVs present in all four enrichment cultures above 0.1% relative abundance

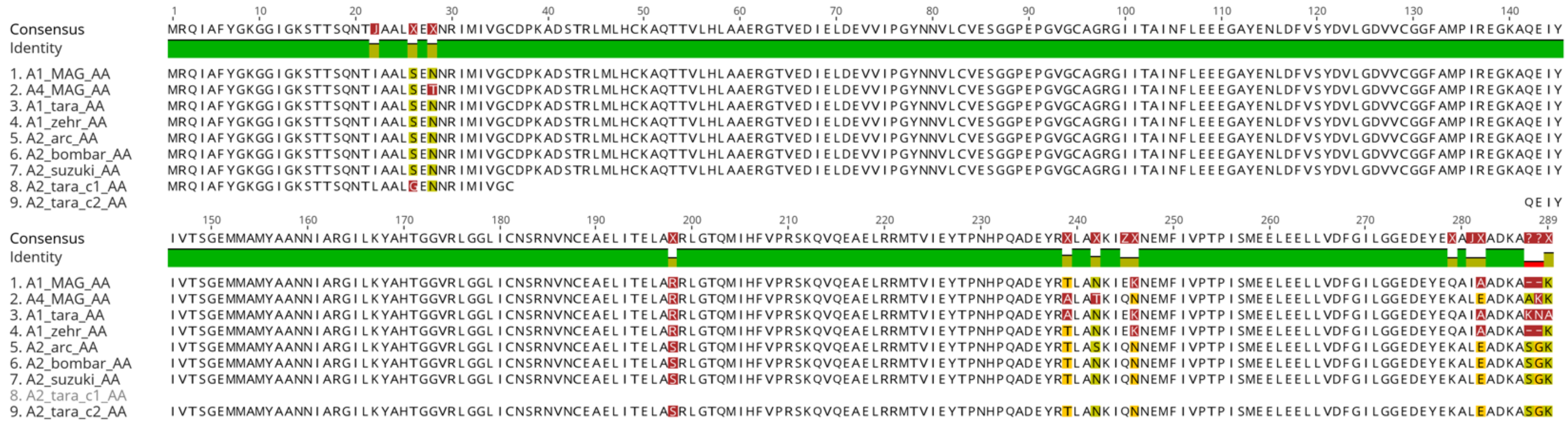

Figure S2. Alignment of NifH amino acid sequences from each of the genomes where *nifH* gene was present.

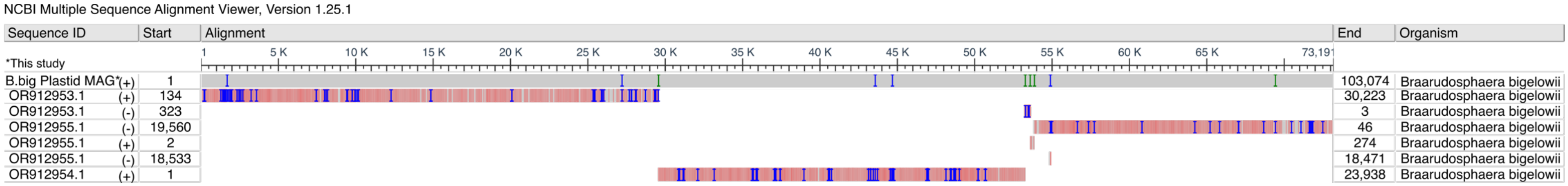


Figure S3. Alignment of our *B. bigelowii* plastid sequence with published *B. bigelowii* chloroplast sequence from Coale et al. (13) (acc: OR912955.1, OR912954.1, OR912953.1).
